# Extreme genome reduction selectively retains modular regulatory architecture in *Prochlorococcus* MED4: conserved transcriptional modules reveal core physiological regulatory programs

**DOI:** 10.64898/2026.04.15.718746

**Authors:** Zachary Johnson, Natalie C. Sadler, Marci R. Garcia, Xiaolu Li, Jordan Rozum, Lindsey Anderson, Tong Zhang, Song Feng, Wei-Jun Qian, Margaret S. Cheung, Pavlo Bohutskyi

## Abstract

*Prochlorococcus* MED4 is a minimal photoautotroph whose extreme genome streamlining extends to its transcriptional regulatory architecture, yet it dominates high-light oligotrophic surface waters and drives marine carbon cycling. Despite ecological significance, MED4 remains genetically intractable, lacking molecular tools to characterize regulatory mechanisms and construct a transcriptome-wide regulatory map. To address this, we assembled an RNA-seq compendium of 253 samples, including 207 new samples capturing transcriptional responses across three classes of experiments: diverse environmental perturbations, a 24-hour circadian cycle, and phage infection. Using independent component analysis (ICA) applied to 247 quality-filtered samples, we identify 32 independently regulated gene set modules in MED4 (iModulons). By comparison, we previously identified 78 iModulons in the model cyanobacterium *Synechococcus elongatus* PCC 7942, revealing how dramatically genome reduction has simplified MED4’s regulatory architecture. Of the 32 iModulons in MED4, 13 are conserved modules that correlate with experimentally validated transcriptional regulons in PCC 7942, identifying regulatory programs that resisted elimination under extreme selective pressure. These conserved modules reveal regulatory programs governing photosynthesis and light responses (RpaB), circadian rhythms (RpaA), and nutrient assimilation (NtcA, PhoB). Known regulator-specific DNA-binding motifs upstream of genes in conserved modules independently support their identification as regulatory targets. Notably, RpaA governs circadian rhythms through three temporally distinct modules in MED4 versus one in PCC 7942, and the RpaB photoprotection module similarly splits into two. This work uncovers the minimal regulatory core governing photosynthesis, circadian rhythms, and C/N/P metabolism in a globally critical but genetically intractable photoautotroph. This approach offers a generalizable framework for regulatory inference beyond model organisms.

**Importance:** The minimal photoautotroph *Prochlorococcus* MED4 possesses only 28 transcriptional regulators, *versus* 150+ found in most cyanobacteria, reflecting a genome streamlined by billions of years of natural selection. This streamlining minimized metabolic costs, enabling MED4 to dominate nutrient-depleted oceans today. This raises a fundamental question: which regulatory programs did nature choose to keep? The answer matters, but MED4 cannot be studied by conventional genetics, placing its molecular machinery beyond direct experimental reach. Instead, we use large-scale computational methods to define groups of co-regulated genes in MED4. By comparing MED4 with a genetically tractable model cyanobacterium, we can distinguish which regulatory programs nature preserved from those it discarded. This work reveals the minimal regulatory architecture sufficient to sustain the smallest oxygenic photoautotroph on Earth. The principles uncovered here, distinguishing essential from dispensable regulatory programs in a naturally streamlined organism, inform the design of minimal photosynthetic platforms for biotechnology.

## 1. Introduction

*Prochlorococcus* is the most abundant photosynthetic organism in the ocean and a cornerstone of marine primary production. Together with *Synechococcus*, these picocyanobacteria are estimated to contribute ∼25% of global oceanic primary production (1). *Prochlorococcus* is distributed throughout tropical and subtropical oligotrophic oceans, occupying diverse light and nutrient niches through genetic diversification across ecotypes (2). This ecological success is linked to extreme genome streamlining, particularly in lineages adapted to high-light (3). The MED4 high-light-adapted ecotype, with one of the smallest genomes of any free-living oxygenic photoautotroph, encodes ∼1800 proteins in a compact 1.66 Mb chromosome, optimized for efficient resource use in nutrient-poor environments (3).

MED4 thrives in surface waters characterized by intense irradiance, oxidative stress, and fluctuating nutrient availability (4). Yet despite these environmental challenges, MED4 encodes a remarkably small repertoire of transcriptional regulators compared to other cyanobacteria (5), including a limited set of sigma factors, two-component systems (RpaA (6), RpaB (7), PhoB (8)), Crp-family regulators (NtcA (9), RbcR (10)), Fur-family metalloregulators (11), and the DNA damage SOS regulator LexA (12). How this minimal regulatory architecture supports robust acclimation across fluctuating light, oxidative, and nutrient conditions remains an open question.

A major barrier to resolving transcriptional mechanisms in *Prochlorococcus* is its genetic intractability (13). While new high-throughput methods are being developed for in vitro characterization of TF-DNA binding (14–17), much of our understanding of transcriptional control in MED4 remains indirect, inferred from computational comparative genomics studies (8, 12, 18) and limited perturbation experiments (19). No systematic transcriptome-wide regulatory map has been constructed for MED4, leaving the relationship between its minimal regulatory architecture and transcriptional responses to environmental conditions largely uncharacterized. This gap highlights the need for alternative strategies that can extract regulatory insights directly from observational data, without relying on extensive genetic manipulation.

Independent component analysis (ICA) provides a data-driven route to overcome these limitations, decomposing transcriptomic expression data into independently modulated gene sets (iModulons) that correspond to underlying regulatory programs (20, 21). This method has been successfully employed to reconstruct known regulons in *Escherichia coli* (20), *Synechococcus elongatus* PCC 7942 (PCC 7942) (22), and emerging model prokaryotic systems (23). Here, we apply this framework to the minimal photoautotroph MED4 to ask which regulatory programs persist when selective pressure has eliminated everything dispensable. Applied to 247 RNA-seq samples covering nutrient and oxidative stress, phage infection, salinity, circadian, and light perturbation conditions, ICA identified 32 iModulons that explain 71.3% of the variance across the gene expression dataset. Comparing regulatory modules with experimentally validated regulons in PCC 7942 reveals how conserved transcriptional architecture controls gene expression in the genetically intractable photoautotroph MED4.

## 2. Results

### 2.1. A condition-diverse RNA-sequencing dataset enables transcriptional architecture discovery in MED4

To characterize transcriptional regulatory control in MED4, we assembled a compendium of 253 RNA-seq samples, including 207 newly collected samples from three experimental projects and 46 retrieved from the public NCBI SRA database (24–26). The newly collected samples include a perturbation dataset, a circadian time-course dataset, and a P-SSP7 phage infection dataset. After quality control (QC, **Methods 5.3**), 247 samples were retained (**Table 1**).

**Table 1.** RNA-sequencing datasets in iModulon training data by project. Samples (n / N) denote the number of samples retained after QC filtering (n) from the initial total (N).

| Dataset | Samples (n/N) | Experimental Conditions | Ref. |
| --- | --- | --- | --- |
| Perturbation | (48 / 52) | Nutrient depletion (– N / P), trace element limitation, high-light exposure, temperature shift, salinity, oxidative stress (H<sub>2</sub>O<sub>2</sub>, metal stress), pH variation, yeast extract and vitamin availability | new |
| Circadian | (42 / 44) | 24-hour circadian cycle (2 h sampling frequency, 12 timepoints, 3 biological replicates); light-decoupling subset at ZT 2, 4 h (extended darkness) and ZT 14, 16 h (extended light) | new |
| P-SSP7 LD | (111 / 111) | P-SSP7 infection experiments, (8-hours, 6 timepoints)<br>Light, high light, dark infected time-series<br>Light & high light uninfected control time-series | new |
| P-HM2 LD | (40 / 40) | P-HM2 infection experiments, (8-hour, 4 timepoints) | (25) |
| Salinity | (6 / 6) | High salinity vs control salinity | (26) |
| **Total** | **(247 / 253)** | **5 datasets; 26 distinct experimental conditions** | — |

new = RNA-sequencing data newly collected and first reported in this study

### 2.2. optICA identifies biologically meaningful transcriptional modules representing core metabolic and stress functions in MED4

iModulons are identified in MED4 using optimal-dimensionality independent component analysis (optICA), a top-performing module detection benchmarked against experimentally validated transcriptional regulons (21). Applied to the 247-sample expression matrix, optICA decomposes the matrix (**X**) into iModulon gene coefficients (**S**) and iModulon activities across experimental conditions (**A**) (**Figure 1a**) (20, 21). Optimal dimensionality, which determines the number of iModulons, is defined by iteratively computing ICA across multiple dimensions and selecting the highest dimensionality that yields the fewest single-gene iModulons (27). In MED4, this approach identifies 32 iModulons that together explained 71.3% of the variance in the gene expression dataset (**Figure 1b** and **Supplemental Data 2**).

**Figure 1.**
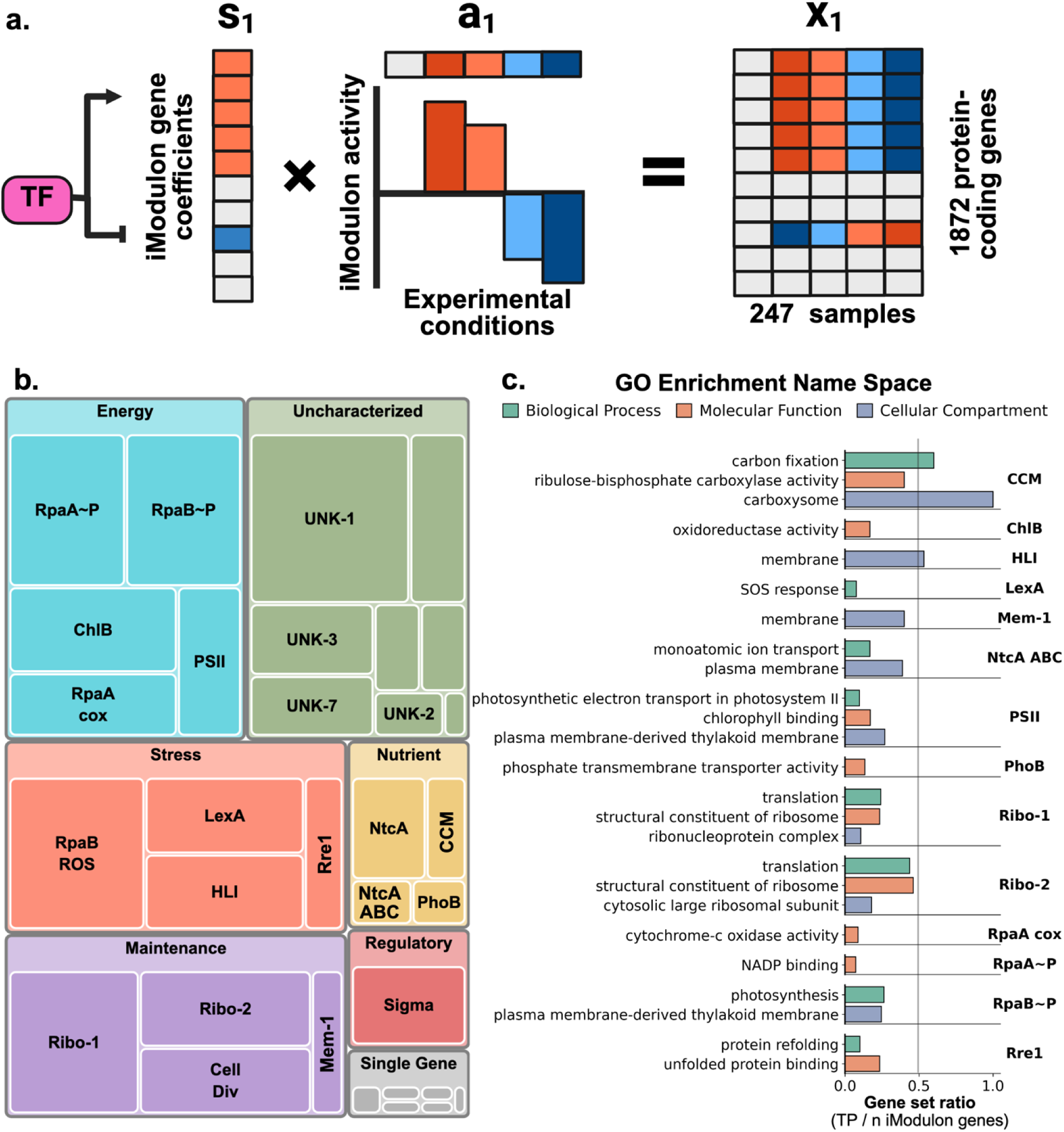
Independent component analysis (iModulons) in MED4 identifies functionally enriched gene sets related to diverse metabolic functions. **a).** Independent component analysis (ICA) decomposes the gene expression matrix (X) into a matrix of gene coefficients across *n* iModulons (S; s_1_…s_n_) and a matrix of iModulon activities across experimental conditions (A; a_1_…a_n_). The product of S and A approximates X, and the explained variance (71.3%) indicates how well this reconstruction captures the original gene expression matrix. b). MED4 iModulons are associated with energy, carbon, nitrogen, and phosphorus metabolism as well as ribosome, protein chaperone, and stress response regulation. The size of each square in the treemap denotes the variance explained by the corresponding iModulon c). GO enrichment analysis (q-value < 0.05) of MED4 iModulon gene sets grouped by GO namespace.

To define iModulon gene sets, k-means clustering was applied to the absolute gene coefficients for each iModulon column of the S matrix to identify genes with significant coefficients (**Methods 5.5**) (20). Of the 32 iModulon gene sets identified, 16 are functionally enriched (q-value < 0.05) with biological processes related to energy metabolism, central nutrient assimilation pathways, and stress responses (**Methods 5.5, Supplemental Data 3**). Energy metabolism iModulons include photosynthesis (RpaB∼P, PSII, ChlB), oxidative phosphorylation (RpaA cox, Ribo-2), and redox homeostasis (RpaA∼P). Nutrient assimilation iModulons cover carbon (CCM, RpaA∼P), nitrogen (NtcA, NtcA ABC), and phosphorus (PhoB) acquisition. Stress-responsive iModulons encompass photoprotection (HLI, RpaB ROS), proteostasis (Rre1), and the DNA damage SOS response (LexA) (**Figure 1b, c**).

### 2.3. Core transcriptional modules are conserved in MED4 despite the reduction in its regulatory architecture due to genome streamlining

Conserved regulons in MED4 were identified using three independent approaches: comparing iModulon gene sets between MED4 and the model cyanobacterium PCC 7942 (**Methods 5.5**), identifying conserved TF DNA-binding motifs upstream of iModulon genes (**Methods 5.6**), and comparing iModulon activity profiles across biologically relevant conditions. To identify regulatory modules conserved between MED4 and PCC 7942 (**Supplemental Data 4**), we compare iModulon gene coefficients (S matrix) across bidirectionally best hit (BBH) orthologs (**Methods 5.6, Supplemental Data 5**) (28).

Of the 32 iModulons in MED4 and 78 in PCC 7942, 13 MED4 iModulons are significantly correlated with a PCC 7942 counterpart (**Supplemental Figure 1, Supplemental Table 1**). Pearson correlation of iModulon gene coefficients across BBH orthologs, with a threshold of r > 0.20, a deliberately inclusive criterion reflecting the expectation that extensive genome reduction and gene content divergence between these phylogenetically distant cyanobacteria will attenuate cross-species correlation even for genuinely conserved regulatory programs. Therefore, motif-based evidence independently validates regulatory conservation (**Table 2**).

**Table 2.** Independent components (iModulons) enriched for known TF DNA-binding motifs in Prochlorococcus marinus MED4 correlate with iModulons enriched for experimentally validated TF-gene binding sites in *Synechococcus elongatus* sp. PCC 7942.

| TF | MED4 iModulon | MED4 TF precision motif (n/N)<sup>a</sup> | PCC 7942 iModulon | PCC 7942 TF precision regulon (n/N)<sup>b</sup> | MED4-PCC7942 iModulon Correlation (Pearson-r)<sup>c</sup> | BBH genes (n) | BBH gene names (MED4) |
| --- | --- | --- | --- | --- | --- | --- | --- |
| **RpaB** | RpaB~P | 0.07 (4/57) | RpaB~P | 0.55 (21/38) | 0.46 | 8 | *psaC, psaA, psaJ, psaK, psbO, psaF, psaD, cpeB* |
|  | RpaB ROS | 0.03 (2/71) | RpaB ROS | 0.59 (17/29) | 0.24 | 5 | *ftsH3, psbD2, hepB, PMM0921, PMM0680* |
|  | Hli | 0.80 (12/15) | RpaB ROS | 0.59 (17/29) | 0.31 | 2 | *rpoD3, hli17* |
| **RpaA** | RpaA~P | 0.10 (7/70) | RpaA~P | 0.66 (60/91) | 0.24 | 9 | *pntA1, zwf, gndA, rpoD6, tal, opcA, pntB, glgP, PMM0383* |
|  | RpaA cox | 0.26 (9/34) | RpaA TOx | 0.70 (14/20) | 0.33 | 5 | *ctaC, ctaB, ctaE, ctaA, ctaD* |
|  | Sigma | 0.79 (11/14) | – | – | – | – | – |
| **NtcA** | NtcA ABC | 0.39 (7/18) | NtcA-2 | 0.79 (15/19) | 0.34 | 5 | *cynB, cynA, cynS, cynD, glnA* |
|  | NtcA | 0.52 (11/21) | NtcA-1 | 0.69 (25/36) | 0.20 | 3 | *ntcA, glnB, glnA* |
| **Rre1** | Rre1 | – | Rre1 HrcA | 1.0 (5/5) | 0.49 | 4 | *groES, dnaK2, groEL1, htpG* |
| **PhoB** | PhoB | 1.0 (15/15) | SphR | 0.12 (2/17) | 0.21 | 2 | *pstC, phoB / sphR* |
| **RbcR<sup>d</sup>** | CCM | – | CCM-1 | – | 0.45 | 3 | *csoS1, cbbS, cbbL* |
| **LexA** | LexA | 0.20 (11/56) | – | – | – | – | – |

**a.** FIMO motif sites identified in the promoter region upstream of operon start site (300bp, q-value < 0.05) **Supplemental Data 6** **b.** iModulon gene set regulatory evidence in *Synechococcus elongatus* PCC 7942 is given in **Supplemental Data 4** [27]. **c.** Bidirectional best hit (BBH) and iModulon correlation between MED4 and PCC 7942 is given in **Supplemental Data 5** [27]. **d.** The RbcR regulon is inferred from the regulatory architecture described for PCC 6803 [8]

Among these 13 correlated iModulons, conserved regulons are identified for six TFs: RpaA, governing circadian regulation; RpaB and Rre1, involved in light stress and proteostasis responses; and PhoB, NtcA, and CCM, regulating phosphorus, nitrogen, and carbon assimilation. One additional iModulon—Sigma—is enriched for RpaA DNA-binding motifs but has no correlated counterpart in PCC 7942. The remaining 19 MED4 iModulons have no correlated counterpart in PCC 7942, likely reflecting regulatory programs unique to MED4’s ecological niche or too diverged in gene content to meet the correlation threshold.

### 2.4. Transcriptional regulation of nitrogen, phosphorus, and carbon assimilation

To characterize how MED4 coordinates nutrient acquisition under resource-scarce conditions, we analyzed transcriptional responses across nitrogen and phosphate depletion, with salinity, temperature, and redox perturbations included as broader environmental context. Three distinct iModulons—NtcA, PhoB, and CCM—correspond to conserved regulators of nitrogen, phosphorus, and carbon fixation, share conserved gene content with PCC 7942 counterparts, with NtcA and PhoB DNA-binding motifs identified upstream of genes in their respective iModulons (**Table 2, Supplemental Data 6**). The condition-specific activation patterns of these iModulons reveal how conserved regulatory circuits are selectively deployed to manage nitrogen, phosphorus, and carbon availability across distinct environmental challenges.

NtcA, the global regulator of nitrogen assimilation, is represented by two iModulons in MED4 and correlates with three iModulons enriched in validated NtcA regulatory targets in PCC 7942 (**Table 2**, **Figure 2a, Supplemental Figure 1**). These iModulons include conserved genes shared with the PCC 7942 NtcA regulon, including the nitrate/cyanate transporter (*nrtABCDS*), the NtcA transcriptional regulator (*ntcA*), and the PII protein regulator (*glnB*). Across both iModulons, 18 genes in 13 operons contain a predicted NtcA-binding site in their promoter regions.

**Figure 2.**
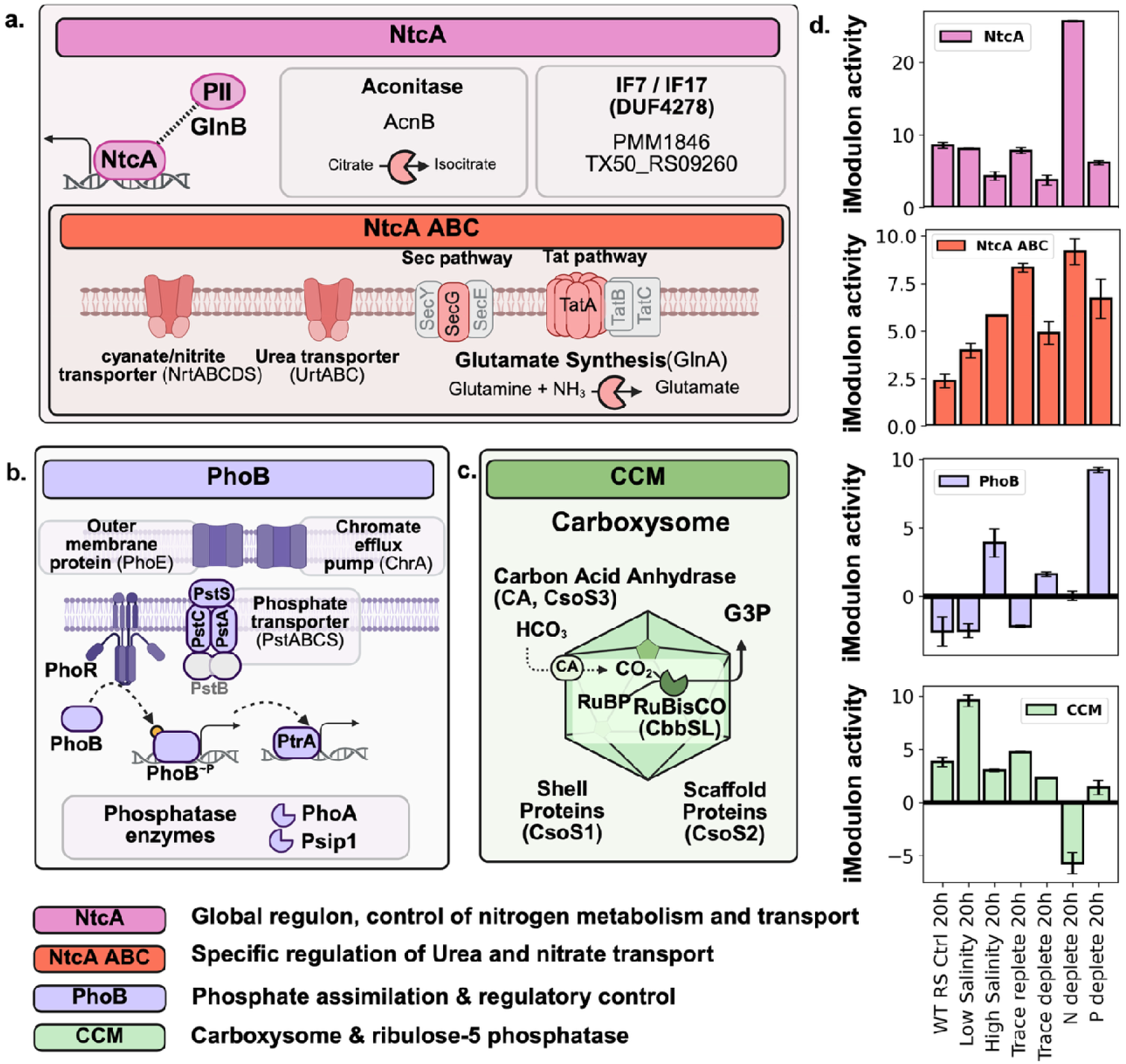
Identification of iModulons controlling nutrient assimilation and iModulon activity responses to environmental perturbations. **a).** Two iModulon gene sets control nitrogen assimilation. iModulon NtcA controls the global nitrogen regulon, including the NtcA-PII regulatory proteins (*ntcA*, *glnB*). NtcA ABC controls nitrate/cyanate and urea assimilation ABC transporters, protein exporter components (*secG*, tatA), and glutamate synthase. **b).** The PhoB iModulon controls phosphate assimilation regulators (*phoBR*, *ptrA*), phosphate transporters (*pstACS, phoE*), and phosphatase enzymes (*psip1*, *phoA*). **c).** The CCM iModulon is correlated with the CCM iModulon in PCC 7942. This iModulon captures the expression of the carboxysome shell proteins (c*soS1-3*) and Rubisco (*cbbSL*) and is likely regulated by RbcR. **d).** iModulon activity responses to environmental perturbations. Conditions include a nutrient-replete control (WT RS Crtl.), salinity perturbations (high/low), trace metal perturbations (replete/deplete), and N/P deplete conditions.

The larger NtcA iModulon (21 genes) is strongly activated in MED4 under nitrogen deprivation (**Figure 2d**). The iModulon includes *ntcA*, glutamate synthase (*glnA*), *glnB*, IF7/IF17 glutamate synthase regulator homologs (PMM1846, TX50_RS09260) (29), ABC transporter components, and other uncharacterized genes with predicted NtcA-binding motifs (**Figure 2a**) (18). The smaller NtcA-ABC iModulon (18 genes) captures expression of the nitrate/cyanate transporter (*nrtABCDS*) and the urea transporter (*urtABC*; **Figure 2a**), as well as membrane transport complex proteins (*secG*, *tatA*). Notably, this iModulon was broadly upregulated across most perturbations (**Figure 2d**), suggesting that these uptake systems respond to environmental cues beyond nitrogen stress.

The PhoB iModulon has the highest coverage of regulator DNA-binding sites, with predicted motifs in the promoter regions of all 15 genes across 13 operons. The iModulon captures the canonical phosphate acquisition system in MED4 and shares conserved genes with the SphR regulon in PCC 7942, including the two-component response regulator (*phoB*/*sphR*) (**Table 2**, **Figure 2b**). This gene set also includes the secondary regulator (*ptrA)*, phosphatases (*phoA*, *psip1*), the outer membrane porin (*phoE*), and the chromate transporter (*chrA*) (8). The iModulon is strongly activated under phosphate-deplete conditions, consistent with the expected PhoB response, which mediates sensing, acquisition, and secondary regulatory control of phosphate metabolism (**Figure 2d**).

Despite its simplified carbon-concentration machinery (CCM) (30), MED4 retains the CCM regulator RbcR (10), which is inferred to regulate the CCM iModulon, a small gene set encoded within a single genomic region and comprising carboxysome shell proteins, carbonic anhydrase, and Rubisco (*cso1–3*, *rbcLS*; **Figure 2c**). A strong correlation between the PCC 7942 and MED4 CCM iModulons (r = 0.45) supports regulatory conservation (**Table 2**). The CCM iModulon is activated under low-salinity conditions and repressed during nitrogen depletion, suggesting coordinated modulation of carbon fixation capacity with broader metabolic state (**Figure 2c)**.

### 2.5. Circadian transcriptional programs are preserved despite a simplified oscillator architecture

Having established the nutrient-responsive regulatory modules, we next characterized iModulon dynamics across a 24-hour diurnal cycle to resolve how MED4 coordinates circadian transcriptional programs despite its simplified oscillator architecture. Circadian regulation coordinates transcriptional programs with daily light–dark cycles in cyanobacteria (31). While canonical circadian control is mediated by the KaiABC oscillator (32), MED4 encodes a simplified circadian architecture lacking KaiA and several input proteins (33). Despite this simplification, diel transcriptional outputs are conserved in MED4 (34). To characterize circadian gene regulation in MED4, cultures were sampled across a 24-hour light–dark cycle, with the first time point collected at dawn immediately prior to lights-on (0 h Zeitgeber time, ZT) (**Figure 3a**). Circadian rhythmicity was assessed using cosinor regression applied to both individual gene expression profiles and iModulon activities (**Methods 5.8, Supplemental Data 7)**.

**Figure 3.**
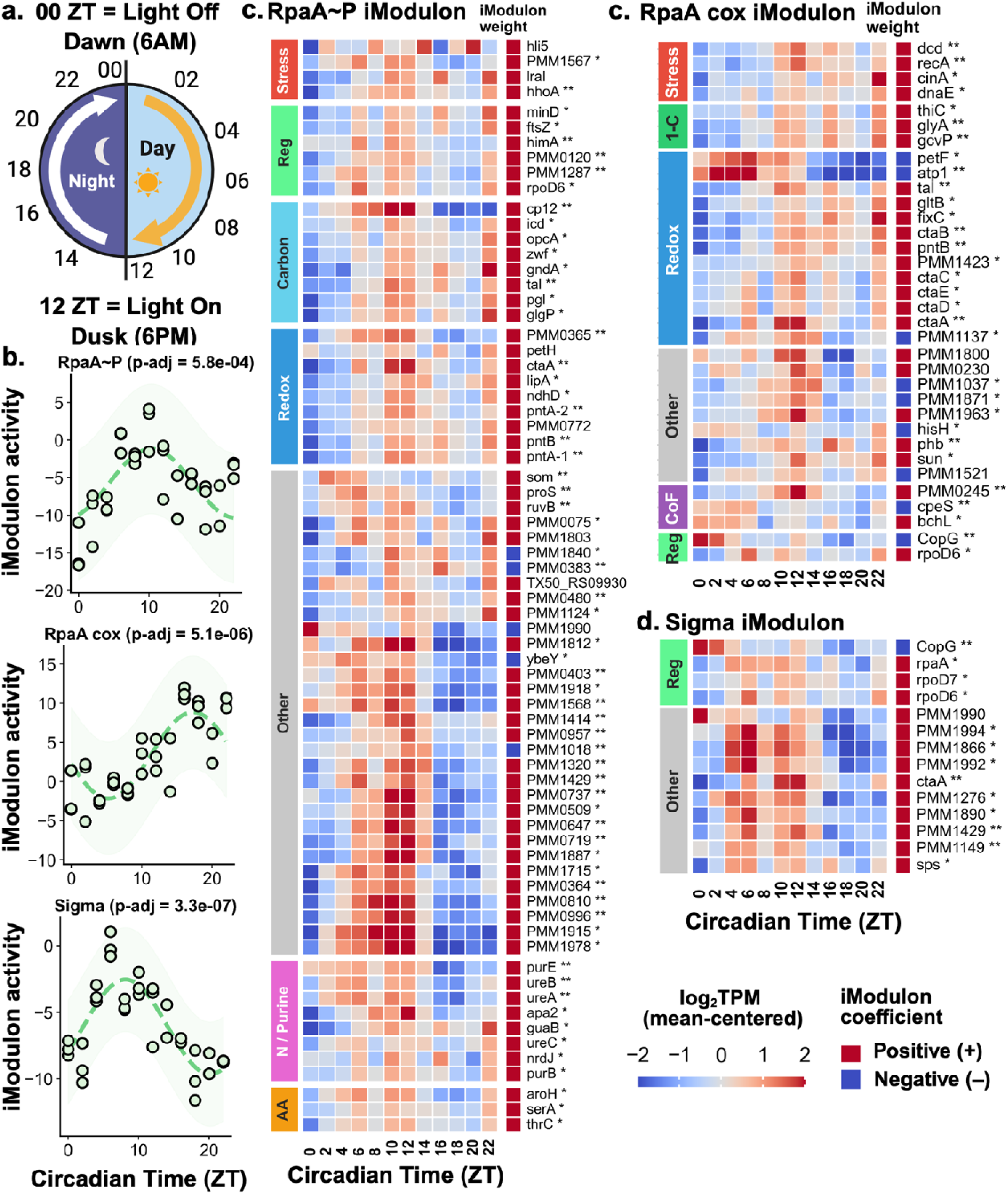
Circadian regulation of iModulons linked to the global circadian regulator RpaA. **a).** Experimental setup for the circadian experiment, the first sample was taken at dawn (ZT = 0, 6 AM) immediately prior to lights turning on. Cells were grown in constant light, then switched to the dark after 12 hours. The dusk time point (ZT=12, 6 PM) was sampled immediately before the lights were turned off. **b).** iModulon activity fit by Cosinor regression for RpaA-associated iModulons **c).** Mean-centered log_2_TPM gene expression for genes identified in the RpaA∼P iModulon clustered by gene set function. Color coding indicates if a gene’s iModulon coefficient is **positively weighted (+, red)**, or **negatively weighted (–, blue)**. Asterisks next to a gene’s name denote whether the gene is identified by a single experimental method (*), or both Cosinor regression (this study) and Fourier analysis on microarray data (**) (34). Genes are linked to stress response (**Stress**), regulatory control of transcription and the cell cycle (**Reg**), central carbon (**Carbon**), redox homeostasis (**Redox**), purine metabolism (**N / Purine**), and amino acid biosynthesis (**AA**). **d).** Mean-centered log2TPM gene expression for genes in the RpaA cox iModulon. The gene set controls the cox operon (oxidative phosphorylation) and redox homeostasis (**Redox**), tetrahydrofolate 1-carbon metabolism (**1-C**), cofactor metabolism (**CoF**), DNA damage repair (**Stress**), and transcriptional regulators; *rpoD6* and CopG family protein (PMM1176) (**Reg**). **e).** Mean-centered log_2_TPM gene expression for genes in the Sigma iModulon. The iModulon gene set includes transcriptional regulators *rpaA*, *rpoD6*, rpoD7/*sigD*, and CopG family protein (PMM1176) (**Reg**).

At the gene level, 1,055 of 1,872 genes showed a significant fit to the circadian model (q-value < 0.01), of which 559 overlapped with previously reported circadian genes in MED4, demonstrating strong agreement with prior microarray-based studies (34). At the iModulon level, 15 of 32 iModulons showed significant circadian rhythmicity (q-value < 1e^−3^, **Supplemental Figure 2, Supplemental Table 2**), with 7 functionally enriched (**Table 3**). The most strongly fit was PhoB (q-value = 2.5 × 10⁻¹²), peaking during the circadian night (∼ 16h ZT). Across significantly rhythmic iModulons, all peaked at dusk or during the circadian night, except for the two associated with ribosome biogenesis (Ribo-1, Ribo-2), which peaked at 2 and 4 h ZT, respectively. Notably, three iModulons associated with the global circadian response regulator RpaA were significantly rhythmic, highlighting its central role in coordinating circadian transcriptional programs in MED4.

**Table 3.**
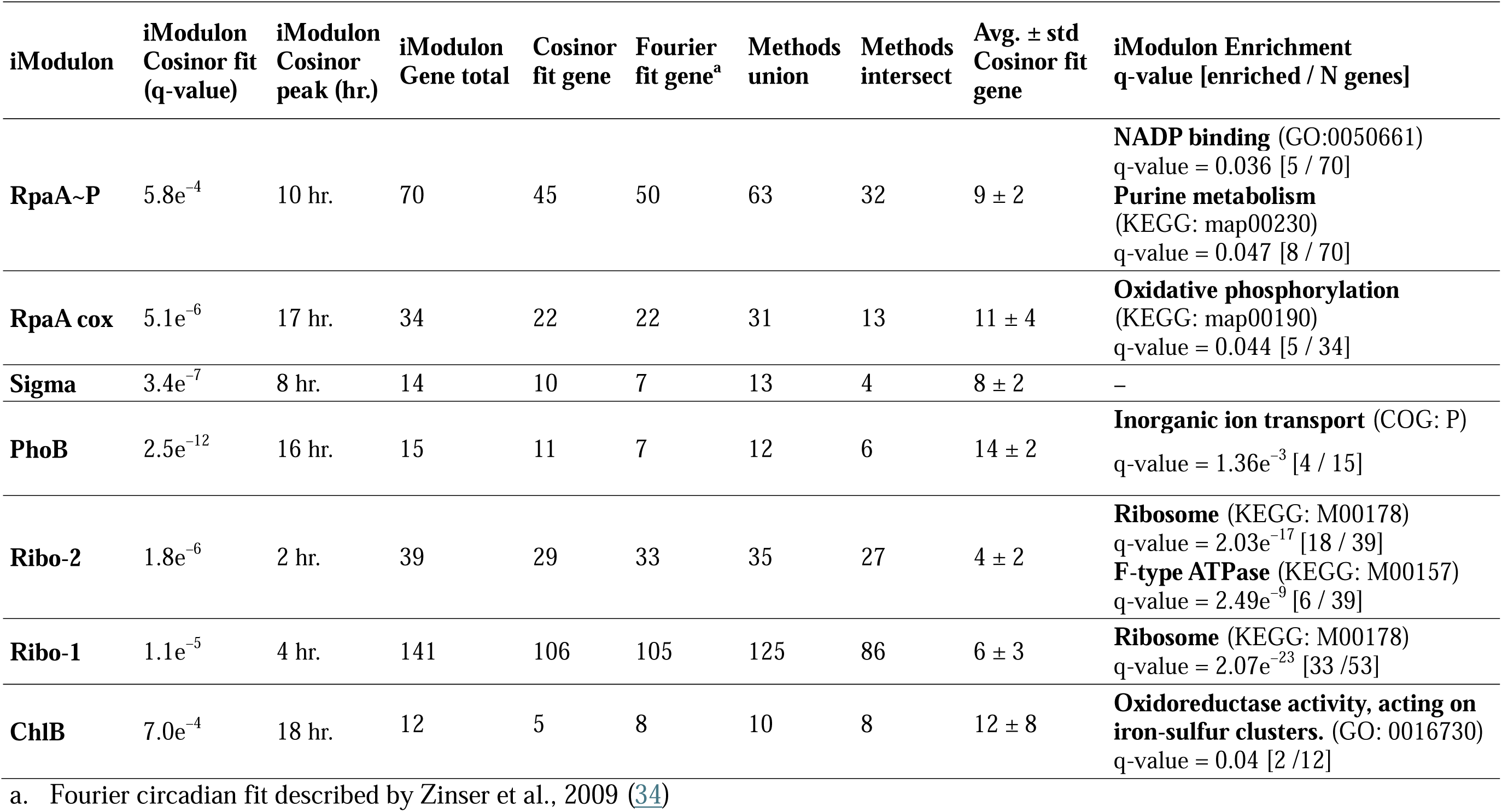
iModulons affected by circadian and light-dependent regulation, excluding uncharacterized and single gene iModulons.

| iModulon | iModulon<br>Cosinor fit<br>(q-value) | iModulon<br>Cosinor<br>peak (hr.) | iModulon<br>Gene total | Cosinor<br>fit gene | Fourier<br>fit gene<sup>a</sup> | Methods<br>union | Methods<br>intersect | Avg. ± std<br>Cosinor fit<br>gene | iModulon Enrichment<br>q-value [enriched / N genes] |
| --- | --- | --- | --- | --- | --- | --- | --- | --- | --- |
| **RpaA~P** | 5.8e<sup>-4</sup> | 10 hr. | 70 | 45 | 50 | 63 | 32 | 9 ± 2 | **NADP binding** (GO:0050661)<br>q-value = 0.036 [5 / 70]<br>**Purine metabolism**<br>(KEGG: map00230)<br>q-value = 0.047 [8 / 70] |
| **RpaA cox** | 5.1e<sup>-6</sup> | 17 hr. | 34 | 22 | 22 | 31 | 13 | 11 ± 4 | **Oxidative phosphorylation**<br>(KEGG: map00190)<br>q-value = 0.044 [5 / 34] |
| **Sigma** | 3.4e<sup>-7</sup> | 8 hr. | 14 | 10 | 7 | 13 | 4 | 8 ± 2 | – |
| **PhoB** | 2.5e<sup>-12</sup> | 16 hr. | 15 | 11 | 7 | 12 | 6 | 14 ± 2 | **Inorganic ion transport** (COG: P)<br>q-value = 1.36e<sup>-3</sup> [4 / 15] |
| **Ribo-2** | 1.8e<sup>-6</sup> | 2 hr. | 39 | 29 | 33 | 35 | 27 | 4 ± 2 | **Ribosome** (KEGG: M00178)<br>q-value = 2.03e<sup>-17</sup> [18 / 39]<br>**F-type ATPase** (KEGG: M00157)<br>q-value = 2.49e<sup>-9</sup> [6 / 39] |
| **Ribo-1** | 1.1e<sup>-5</sup> | 4 hr. | 141 | 106 | 105 | 125 | 86 | 6 ± 3 | **Ribosome** (KEGG: M00178)<br>q-value = 2.07e<sup>-23</sup> [33 / 53] |
| **ChlB** | 7.0e<sup>-4</sup> | 18 hr. | 12 | 5 | 8 | 10 | 8 | 12 ± 8 | **Oxidoreductase activity, acting on**<br>**iron-sulfur clusters.** (GO: 0016730)<br>q-value = 0.04 [2 / 12] |

a. Fourier circadian fit described by Zinser et al., 2009 (34)

### 2.6. Response regulator RpaA governs circadian transcriptional programs through three distinct iModulons

Of the three RpaA-associated iModulons in MED4, two—RpaA∼P and RpaA cox—show significant gene set correlations with iModulons in PCC 7942, whereas the Sigma iModulon has no PCC 7942 counterpart and represents a regulatory program unique to MED4. Notably, the Sigma iModulon shows the highest coverage of RpaA DNA-binding motif sites upstream of operon transcription start sites (**Supplemental Data 6**).

The RpaA∼P iModulons in MED4 and PCC 7942 are moderately correlated (Pearson-r = 0.24) but capture core regulatory programs required for balancing redox during circadian night. In PCC 7942, the iModulon is independently validated, capturing 66% of RpaA ChIP-seq binding targets (60/91 genes) (6, 35). Mapped BBH orthologs across these transcriptional modules identify conserved regulatory control of glycogen catabolism (*glgP*), NADPH regeneration (*pntAB, zwf*, *opcA*, *gndA*, *tal*), and transcriptional regulation (*rpoD6*). Beyond these shared orthologs, the MED4 iModulon additionally captures known circadian regulatory outputs, including a redox-sensing regulator of the Calvin-Benson Cycle (*cp12*) (36), a nucleoid protein associated with genome supercoiling (*himA*) (37), NDH1 subunit (*ndhD*) (38), and cell division proteins (*ftsZ*, *minD*) (39, 40). The RpaA∼P iModulon activity peaks around 10 h ZT, slightly after the average gene expression peak at ∼9 h ZT, and consistent with reports of maximum RpaA phosphorylation occurring immediately prior to dusk (**Figure 3b, c**) (6).

The RpaA cox iModulon in MED4 is correlated with an RpaA-enriched iModulon regulating expression of terminal oxidase (RpaA TOx) in PCC 7942 (**Table 2**, r = 0.43). In MED4, the iModulon captures transcriptional control of the cytochrome-c oxidase *cox* operon, DNA repair mechanisms, 1-carbon and cofactor metabolism, as well as transcriptional regulators *rpoD6* and the CopG family regulator (PMM1176) (**Figure 3d**). In MED4, iModulon activity peaks around 17 h ZT (Figure 3b), whereas average gene expression peaks at 11 h ZT.

Finally, the Sigma iModulon has no correlated counterpart in PCC 7942 but represents a divergent circadian regulatory program in MED4, which is significantly enriched for RpaA-binding sites upstream of operon transcription start sites (11/14 genes). Here, *rpaA*, *rpoD6*, *rpoD7*/*sigD* (PMM0577), and the CopG family regulator are co-regulated (**Figure 3e**). Gene expression and iModulon activity peak around 8 h ZT (**Figure 3c**), the earliest among RpaA-associated iModulons.

Collectively, the three RpaA-associated iModulons define the core circadian transcriptional landscape in MED4, yet cyanobacteria must also respond dynamically to unexpected changes in light availability, raising the question of how these circadian programs interact with light-dependent regulation.

### 2.7. Light perturbations differentially activate RpaA- and RpaB-associated iModulons

While circadian entrainment coordinates anticipatory transcriptional responses to predictable light–dark cycles, cyanobacteria must also respond rapidly to unexpected changes in light availability (41). To evaluate how light sensing and circadian regulation are coordinated in MED4, transcriptional regulation was assessed under extended darkness during the entrained circadian day and extended light during the entrained circadian night (**Figure 4a**). Differential iModulon activities (DIMAs) and differential gene expression analysis (**Methods 5.8**) were performed to characterize how gene expression responds in the light-perturbed circadian conditions (**Figure 4b, c**) (42, 43).

**Figure 4.**
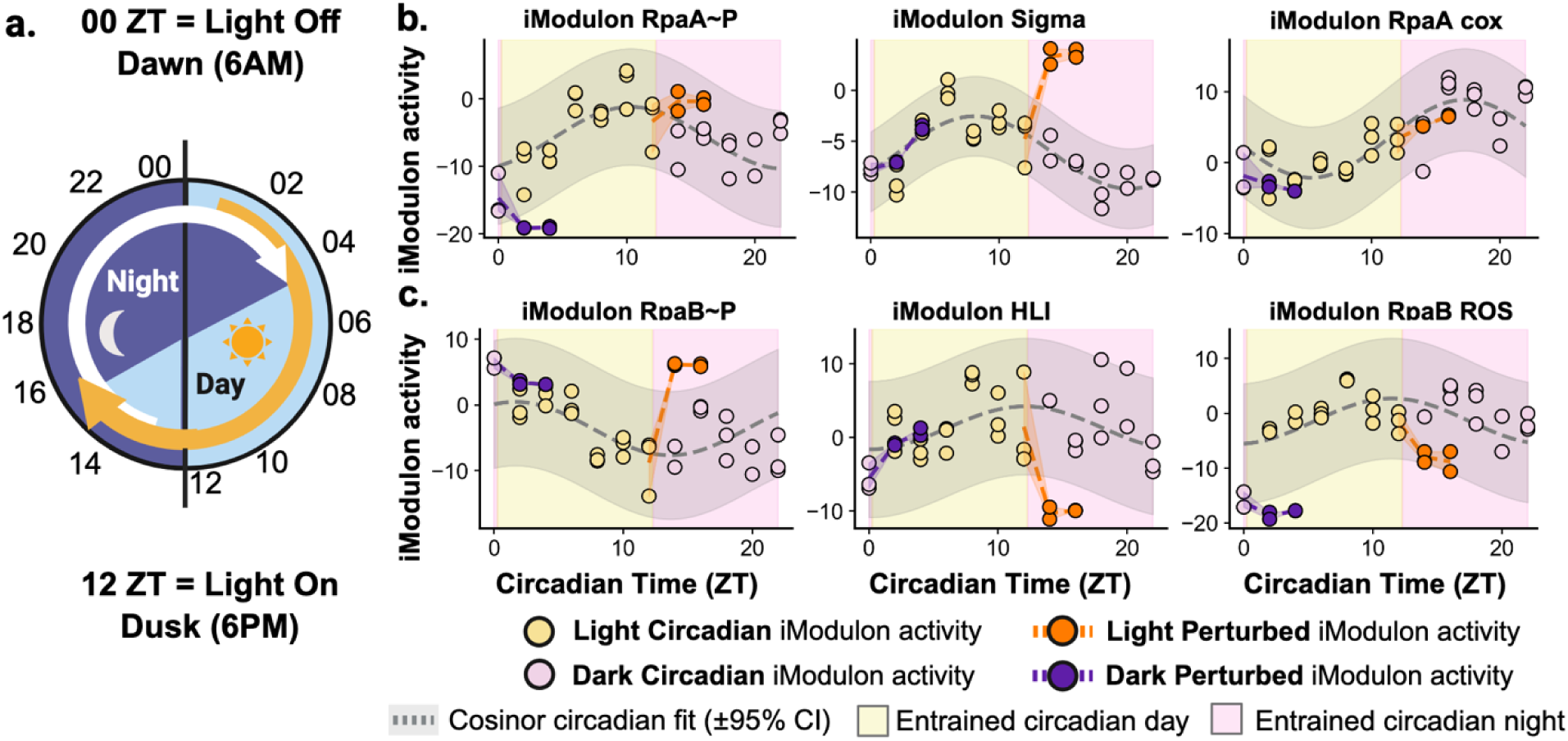
Light-driven transcriptional changes at the transcriptional module level in Prochlorococcus MED4. **a).** Experimental design for the light-perturbed circadian experiment in Prochlorococcus MED4. Additional samples were taken during the circadian transcriptomics experiment for cultures exposed to extended periods of darkness (2, 4 h ZT) after circadian entrained dawn (0 h ZT) and extended periods of light (14, 16 h ZT) after circadian entrained dusk. **b).** RpaA-associated iModulon activity for samples taken during the circadian cycle and during light perturbation. RpaA∼P demonstrates reduced activity during extended darkness during the circadian day. The Sigma iModulon increased activity during the extended light period. The RpaA cox iModulon has no differential activity in response to light. **c).** RpaB-associated iModulon activity in response to light perturbations

Comparison of differential gene expression analysis identified 897 genes differentially expressed during extended darkness in the circadian day, and 121 during extended light in the circadian night (p-adjusted < 0.05; **Supplemental Data 8**). Differential iModulon activity analysis (DIMA, p-adjusted < 0.05) identified 14 iModulons differentially activated by extended darkness and 11 by extended light (**Supplemental Figure 3, Supplemental Tables 3** and **4**). Across iModulons, those associated with RpaA and RpaB were among the most differentially activated. RpaA∼P showed significantly reduced activity during extended darkness, whereas the Sigma iModulon showed significantly increased activity during extended light (**Figure 4b**). All three RpaB-associated iModulons were also differentially activated: RpaB∼P was strongly activated during extended light, while both RpaB ROS and HLI were repressed by extended light (**Figure 4c**). These results demonstrate that both the circadian regulator RpaA and the light-responsive regulator RpaB are acutely sensitive to unexpected changes in light availability, motivating closer characterization of the RpaB gene sets and their role in balancing photosynthetic electron transport.

### 2.8. RpaB iModulons coordinate photosynthesis, photoprotection, and oxidative phosphorylation

In cyanobacteria, RpaB functions as the global regulator of photosynthesis, controlling genes that respond to both low-light and high-light intensities (7, 44, 45). In PCC 7942, five iModulons are significantly enriched in RpaB binding targets (22). Of these, two share homology with three MED4 iModulons. Characterization of these iModulon gene sets demonstrates strong evolutionary conservation of the regulatory control of photosynthesis.

The primary iModulon controlling expression of the photosynthetic protein complexes and pigment genes in MED4 and PCC 7942 is RpaB∼P (r = 0.48, **Table 2**). The MED4 iModulon captures co-regulated expression of photosystem II (PSII), photosystem I (PSI), cytochrome b_6_f (B6f), and light-harvesting proteins (LH) (**Figure 5a, b**). The gene set also identifies co-regulation of pigment biosynthesis, including genes involved in heme biosynthesis (*hemEJ*) and divinyl chlorophyll biosynthesis (*acsF*, *chlLNP*). Of the RpaB∼P iModulon genes, only one gene is involved in carotenoid biosynthesis (*crtL2*). The broader photoprotection program—including carotenoid metabolism, ROS defense, and PSII repair—is controlled by the iModulons RpaB ROS and HLI.

**Figure 5.**
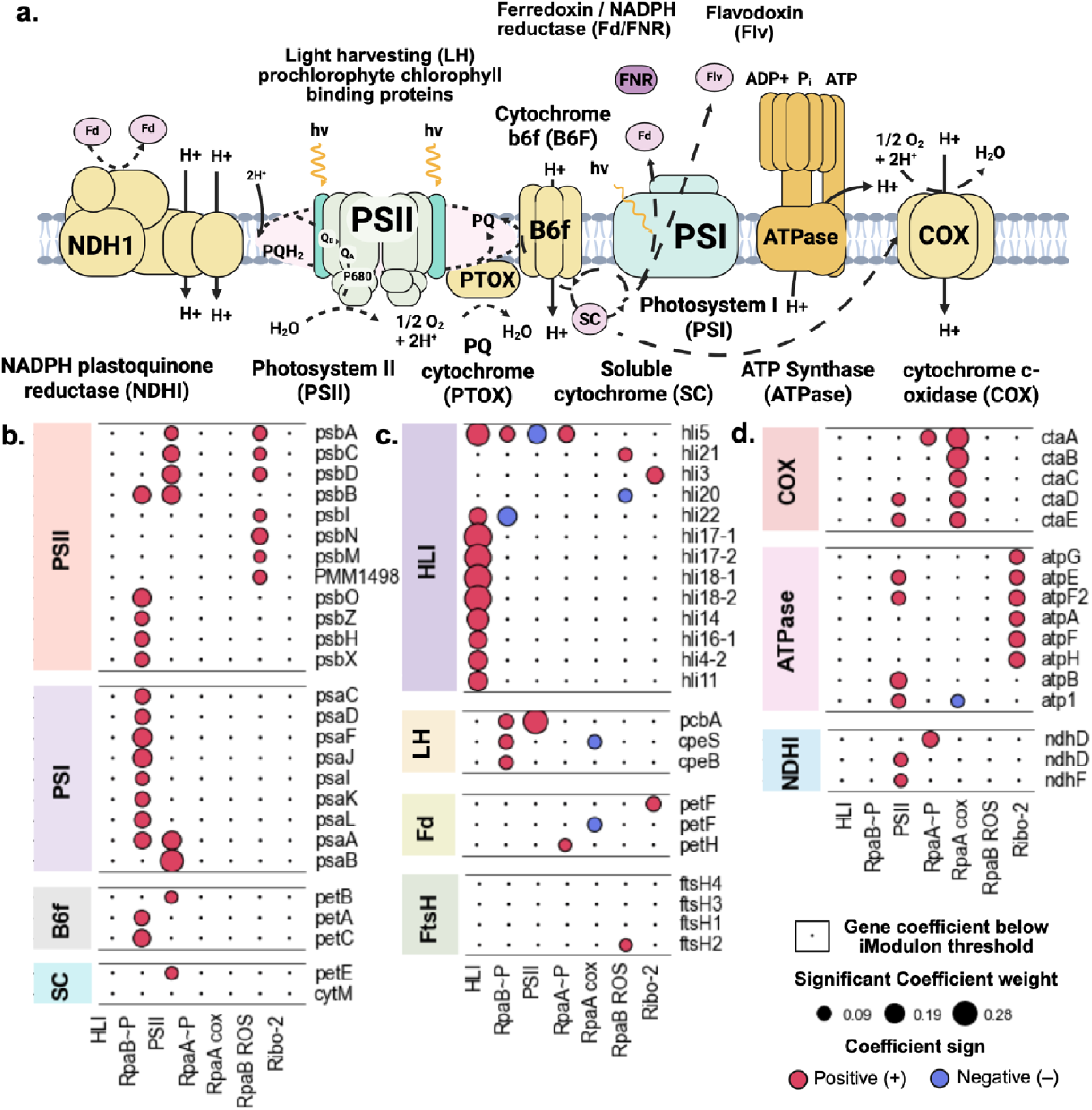
Regulatory control of electron transport protein complexes in Prochlorococcus marinus MED4. **a).** Cartoon depiction of the electron transport chain in MED4 where primary photosynthetic complexes and proteins are highlighted in green, photoprotective mechanisms are highlighted in red, and oxidative phosphorylation pathways in yellow. **b).** iModulons gene coefficients for primary photosynthetic electron transport processes are primarily captured by the iModulons RpaB∼P, PSII, and RpaB ROS. **c).** Photoprotective mechanisms in MED4 are captured by the iModulons RpaB ROS and HLI. **d).** Oxidative phosphorylation pathways are controlled by iModulons RpaA cox, PSII, and Ribo-2 (F-type ATP Synthase).

The RpaB ROS iModulon is one of two MED4 iModulons correlated with the single stress response iModulon in PCC 7942 (RpaB ROS; **Table 2**). It shows moderate correlation (Pearson-r = 0.24) and captures core photosystem proteins, including PSII core subunits, PSII repair and assembly proteins (*psbIMN*), and the PSII high-light adaptation protein PMM1498—a homolog of slr0320—previously shown to be crucial for high-light adaptation in *Synechocystis* PCC 6803 (**Figure 5a**) (46). Additionally, the iModulon captures the expression of *rpaA*, ferredoxin (PMM0316), carotenoid metabolism (*sds*, *zds*, *pds*, PMM0280), and ROS defense genes (*flvB*, *gst1*, *btuE*, *rub*, *ftsH2* PMM1015, PMM1359, PMM0345). The HLI iModulon, the second MED4 iModulon correlated with RpaB ROS in PCC 7942 (Pearson-r = 0.31), captures the expression of high-light inducible (HLI) proteins (**Figure 5b**) and the sigma factor (*rpoD3*), both previously linked to the RpaB high-light response in cyanobacteria (5, 47).

In addition to the RpaB-regulated photosynthetic and photoprotective modules, three iModulons not associated with RpaB coordinate oxidative phosphorylation processes in the electron transport chain. The RpaA cox iModulon controls expression of the cytochrome-c oxidase *cox* operon, linking circadian regulation to respiratory electron transport during circadian night. The PSII iModulon broadly coordinates photosynthetic and oxidative phosphorylation processes, controlled by an as-yet unidentified regulator. Finally, the Ribo-2 iModulon is enriched in both ribosomal proteins and ATPase subunits, suggesting coordinated control of protein biosynthesis and ATP synthesis (**Figure 5d**). Together, these modules demonstrate that oxidative phosphorylation and energy conversion in MED4 are distributed across multiple independently regulated transcriptional programs coordinated by both circadian (RpaA), light-responsive (RpaB), and uncharacterized transcriptional regulators.

## 3. Discussion

By applying ICA to a compendium of 247 RNA-seq expression profiles spanning nutrient limitation, high light and oxidative stress, circadian, phage infection, salinity, and light perturbation conditions, we identify transcriptional modules governing carbon concentration (CCM), nutrient acquisition (NtcA, PhoB), circadian regulation (RpaA), and light- responsive control of photosynthesis (RpaB) in *Prochlorococcus* MED4. Comparative analysis based on bidirectional best hit ortholog mapping and iModulon gene coefficient correlation with *Synechococcus elongatus* PCC 7942 reveals that MED4 encodes just 32 iModulons compared to 78 in PCC 7942, consistent with the dramatic regulatory simplification expected from extreme genome streamlining. Of these, 13 MED4 iModulons are conserved across these phylogenetically distant cyanobacteria, the iModulons regulated by RpaA, RpaB, PhoB, and NtcA, independently supported by TF DNA-binding motifs upstream of iModulon genes. These conserved modules define the core regulatory circuits whose function resisted elimination under the selective pressures driving MED4 genome reduction. The computational framework—combining large-scale transcriptomics, ICA, cross-species comparative analysis, and independent motif validation—demonstrates a generalizable strategy for inferring regulatory control in organisms that resist direct genetic characterization.

### 3.1. Conserved transcriptional regulation of carbon concentration machinery despite distinct CCM architectures

Cyanobacteria require carbon concentrating mechanisms (CCMs) to elevate CO₂ around RuBisCO and suppress photorespiration (48, 49). MED4 and PCC 7942 employ distinct CCM architectures, yet both retain conserved transcriptional regulation of core CCM operons.

*Prochlorococcus* MED4 possesses an α-carboxysome architecture, consisting of a catalytically efficient but low CO₂ specificity Type IA RuBisCO (*cbbLS*), with carboxysome shell proteins (*csoS1,2,4*) and carbonic anhydrase (*csoS3*), all encoded within a single genomic locus (30). In contrast, PCC 7942 encodes a β-carboxysome architecture with CCM components dispersed across multiple operons, using Type IB RuBisCO (*rbcLS*), which has lower catalytic efficiency but higher CO₂ specificity (50). Despite these architectural differences, transcriptional modules are resolved for the core CCM locus in both organisms, capturing RuBisCO, primary shell proteins, and scaffolding proteins, whereas assembly factors, minor shell components, and carbonic anhydrase are controlled independently in a species-specific manner.

Notably, the MED4 iModulon reveals that a subset of the core CCM genomic region is differentially regulated despite genes being co-encoded within the same operon. The iModulon includes *csoS2-3* but excludes *csoS4A* and *csoS4B*, suggesting a potential post-transcriptional control mechanism, such as differential mRNA stability, antisense RNA-mediated protection, or riboswitch activity, leading to differential accumulation of transcripts within a single operon (5, 51, 52).

### 3.2. Conserved and divergent circadian programs coordinated by RpaA and type II sigma factors

Global circadian control in both MED4 and PCC 7942 is coordinated by RpaA (6) through a type II sigma factor cascade (53). In both cyanobacteria, iModulons identify co-regulation of core RpaA programs with type II sigma factor RpoD6, though MED4 utilizes a phylogenetically distinct sigma factor (RpoD7*/*SigD) (34), and lacks homologs of circadian-coordinating sigma factors in PCC 7942 and PCC 6803: RpoD5 (SigC) and RpoD2 (SigB) (53, 54). In MED4, two RpaA-associated iModulons (RpaA∼P and RpaA cox) correlate with iModulons enriched in validated RpaA targets in PCC 7942 (**Table 2**), while a third (Sigma) represents a regulatory program unique to MED4.

Conservation of the RpaA∼P iModulon in both organisms underscores the essentiality of circadian-gated metabolic programs during the light-to-dark transition. The conserved co-regulation of the oxidative pentose phosphate pathway (OxPPP), pyridine nucleotide transhydrogenase (PNT), and glycogen phosphorylase (GlgP) reflects a shared requirement for coordinated glycogen mobilization and NADPH regeneration; disruption of RpaA leads to redox imbalance and a lethal phenotype in PCC 7942 (55). This program is further reflected in MED4, preserving redox regulation of the Calvin-Benson Cycle via CP12 (36), modulation of genome accessibility by HimA (37), and gating of cell division through FtsZ and MinD (39, 40). Similarly, conserved is the regulatory control of the *cox* operon (*ctaABCDE*) in the RpaA cox iModulon, suggesting that the aa₃-type cytochrome-c oxidase represents a core circadian program, while additional terminal oxidases (bd-quinol, cbb3-type) in PCC 7942 may provide metabolic flexibility, but are non-essential for circadian function in MED4.

The Sigma iModulon represents a divergent circadian program in MED4, showing the greatest enrichment of RpaA binding motifs among RpaA-associated iModulons (79% of genes, 11/14), and including downstream regulators *rpaA*, *rpoD6*, and putative CopG family TF (PMM1176), but notably not *rpoD7*/*sigD* (PMM0577). The negative gene coefficients of the putative CopG family transcription factor in both the Sigma and RpaA cox iModulons suggest it may function as an opposing regulatory input modulating its activity (**Figure 3d**).

### 3.3. Conserved RpaB regulatory architecture coordinates photosynthesis and photoprotection across divergent light-harvesting systems

In the absence of the circadian clock protein KaiA, MED4 depends on environmental light signals to entrain its truncated Kai oscillator, placing additional regulatory weight on light-responsive transcriptional programs. Central among these are the genes encoding its simplified light-harvesting complexes, the prochlorophyte chlorophyll-binding proteins (Pcbs), which function with divinyl chlorophyll a and b derivatives (56), rather than the phycobilisome-based antenna systems of PCC 7942. Despite this fundamental architectural difference, the RpaB∼P iModulon reveals that transcriptional coordination of PSI and antenna protein genes is conserved as a single co-regulated unit in both organisms (**Table 2**), demonstrating that conserved regulatory architecture can persist despite divergent evolution of the underlying protein components.

Regulatory control of photoprotection shows greater divergence, with the single RpaB ROS iModulon in PCC 7942 mapping to two distinct modules in MED4: RpaB ROS and HLI. The HLI iModulon shows the greatest enrichment of RpaB binding motifs among RpaB-associated iModulons, whereas neither RpaB∼P nor RpaB ROS shows comparable enrichment, paralleling the concentration of circadian RpaA iModulon motifs within the Sigma iModulons. This divergence likely reflects integration of additional regulatory inputs: the HLI iModulon includes the type II sigma factor *rpoD3*, while the RpaB∼P iModulon includes *rpaA* and a putative CRP-family redox-responsive regulator (PMM0806) (57), the latter also appears in the proteostasis (Rre1) iModulon, positioning it at the interface of photoprotection and protein quality control.

Together, these results suggest that while the core architecture of RpaA/RpaB-mediated circadian and light control is conserved between marine and freshwater cyanobacteria, distinct regulatory layers have emerged independently in each lineage. Candidate regulators in MED4 that potentially contribute to these lineage-specific programs include the CopG and CRP family regulators, and small non-coding RNAs—each warrants further investigation as genetic tools for MED4 continue to develop.

## 4. Conclusion

By applying unsupervised machine learning, specifically ICA, to a compendium of 247 RNA-seq profiles, we constructed the first transcriptome-wide regulatory map for *Prochlorococcus marinus* MED4, an ecologically dominant but genetically intractable photoautotroph. The RNA-seq compendium assembled here is publicly available, providing a community resource for future regulatory studies in this organism with a streamlined genome and minimized regulatory architecture. MED4’s architecture includes just 32 transcriptional modules (iModulons) compared to 78 in the regulatory-replete, genetically tractable freshwater model *Synechococcus elongatus* PCC 7942, quantifying how dramatically genome reduction has simplified regulatory control. Of these, 13 are conserved modules shared between the two organisms, spanning carbon concentration, circadian control, photosynthetic regulation, and nutrient assimilation. The conservation of these regulatory programs despite extensive genome reduction suggests they represent core organizational principles of cyanobacterial gene regulation rather than species-specific adaptations. Strikingly, conserved regulators in MED4 govern broader and more functionally diverse programs than their PCC 7942 counterparts, exemplified by RpaA splitting circadian control across three temporally distinct modules and RpaB photoprotection resolving into two, suggesting regulatory subfunctionalization accompanied genome reduction. These computationally derived modules provide experimentally testable predictions of regulatory structure in MED4, offering a roadmap for prioritizing regulatory targets and accelerating regulatory discoveries as genetic tools for this organism continue to develop.

## 5. Methods

### 5.1. Strain cultivation and adaptation to Salish seawater

*Prochlorococcus marinus* subsp. *pastoris* CMP1986 (MED4) was obtained from the Provasoli-Guillard National Center for Marine Algae and Microbiota (NCMA, East Boothbay, ME). Cyanophage P-SSP7 was provided by Allison Coe (Chisholm Laboratory, MIT). For routine maintenance and small volume batch perturbation experiments, MED4 was cultured in Pro99 medium prepared with Sargasso Seawater (NCMA), filtered through a 0.2-μm filter, and autoclaved at 121°C for 30 minutes. After cooling, sterile seawater was amended with Pro99 nutrients, including nitrogen, phosphorus, and trace metals (58). To enable large-volume experiments, including P-SSP7 phage infection, MED4 was gradually acclimated to local Salish Sea water over a two-month period (59). For optimal growth in Salish Sea water, the medium was supplemented with 2× Pro99 nitrogen and phosphorus while maintaining 1× trace metals, which yielded more stable growth and higher biomass density (59). Cultures were maintained at 21°C under a 12 h light: 12 h dark cycle at ∼45 µmol photons m⁻² s⁻¹.

### 5.2. RNA-seq sample processing

For all newly generated RNA-seq samples (spanning 3 experimental projects, 207 samples total), RNA was extracted from exponentially growing cells using the Qiagen RNeasy Plus Mini Kit according to the manufacturer’s instructions. Samples underwent DNase treatment and rRNA depletion before paired-end sequencing (2 × 150 bp) with single-indexing on the Illumina platform at Azenta US, Inc.

### 5.3. RNA-seq experimental conditions

Three experimental datasets were generated in this study; the sample cultivation conditions described in the manuscript are included below. Detailed descriptions of individual samples and procedures are found in **Supplemental Methods 1 and S. Data 1.**

#### Circadian time-course dataset with light-decoupling subset

To characterize diel transcriptional programs in MED4, cultures entrained to a 12 h light:12 h dark photoperiod for at least 180 days (Methods 5.1) were pooled from four 1 L bottles to ensure homogeneity and distributed into 50 mL vented-cap tubes (30 mL per tube). Samples were collected at 2-hour intervals across a complete 24-hour cycle (12 timepoints), with three biological replicates per timepoint (36 samples total). Each biological replicate was derived from two pooled technical replicate tubes. For RNA-seq, 10 mL of culture was harvested by centrifugation (6,500 × g, 6 min, 4°C), snap-frozen in liquid nitrogen, and stored at −80°C. Dark-phase samples were processed under dim green light to prevent inadvertent light exposure.

To distinguish circadian clock-driven responses from direct light-dependent responses, an additional light-decoupling subset was collected at ZT 2 and 4 h under extended darkness (subjective circadian day) and at ZT 14 and 16 h under extended light (subjective circadian night), using the ZT 0 circadian time-course sample as baseline. This subset was collected as biological duplicates. Complete experimental protocols and sample metadata are provided in Supplemental **Methods S2–S3 and Supplemental Data 1**.

#### Environmental perturbation dataset

To characterize transcriptional responses to diverse environmental stressors, batch perturbation experiments were performed across 26 distinct conditions. Exponentially growing cells were collected from a 2 L photobioreactor maintained at 21°C under ∼45 μmol photons m⁻² s⁻¹, and 20 mL aliquots were distributed into 50 mL bio-reaction tubes with 0.22 µm hydrophobic vented membrane caps (CELLTREAT Scientific Products, #229475), enabling gas exchange under aseptic conditions. Perturbation conditions included nitrogen limitation, phosphate limitation, trace element limitation, high-light exposure (200–315 µmol photons m⁻² s⁻¹), pH variation, temperature shift (26°C), sudden availability of yeast extract or vitamins, salinity change, and oxidative stress (H₂O₂ and metal stress). For nutrient limitation conditions, cells were harvested by centrifugation (4,500 rpm, 10 min), washed in 10 mL of the corresponding depleted medium, and resuspended in 20 mL of fresh depleted medium. Incubation times for nutrient depletion conditions and most environmental perturbations were 20 h; for acute stress conditions—high-light, high-temperature, and oxidative stress (H₂O₂)—incubation was shortened to 4–5 h to capture early transcriptional responses and minimize confounding effects of cell death. Complete details for each condition are provided in **Supplemental Data 1**. Cells were harvested by centrifugation and frozen at −80°C prior to RNA extraction.

#### Public RNA-seq datasets

Two previously published RNA-seq datasets were retrieved from NCBI Sequence Read Archive (SRA) (24) using the Entrez Direct API (60) and incorporated into the compendium. The P-HM2 phage infection dataset (40 samples) comprises 8-hour light-dark infection time-series experiments (25), and the MED4 salinity dataset (6 samples) captures transcriptional responses to high salinity conditions (26). The combined pre-QC dataset of newly generated and publicly available RNA-seq profiles consisted of 253 samples.

### 5.4. RNA-seq processing and quantification

Reads were aligned to the *Prochlorococcus* MED4 RefSeq reference (NC_005072.1; released January 26, 2024) (61). Functional annotation was performed using EggNOG-mapper (v5) (62), retrieving KEGG (63) and COG (64) category assignments. Gene Ontology (65) and additional protein annotation were retrieved from UniProtKB (66). Reads were quality-trimmed with TrimGalore v0.6.10 (67, 68), and quality was assessed with FastQC v0.12.1 (69). Alignment and read quantification were performed with Rsubread v2.20 (70). Alignment and quantification quality was performed with Samtools v1.21 (71), Qualimap v2.3 (72), and MultiQC v1.28 (73).

### 5.5. RNA-seq QC and computation of iModulons

Sample QC was adapted from the *iModulonMiner* workflow (42, 74). Genes were retained if their CDS length exceeded 100 bp and the median count exceeded five. Samples were retained based on total CDS counts (CDS > 200,000) and biological replicate correlation (Pearson-r > 0.90); samples identified as outliers during global sample clustering were dropped (**Supplemental Methods 2.1, Supplemental Table 5**). After quality filtering, 247 of 253 samples were retained and used for computational analysis. Expression values were transformed to log_2_TPM and centered on reference conditions (**Supplemental Methods 1, Supplemental Data 1)**. Optimal-dimensionality independent components were computed using optICA (27, 42), and a k-means threshold with manual adjustment was used to define gene inclusion in an iModulon gene set (**Supplemental Methods 2.2, Supplemental Figure 6**).

### 5.6. Comparative analysis of transcriptional modules

Comparative analysis was performed following the pymodulon workflow (42). BBH orthologs were identified using BLAST (75), filtering out hits with an e-value < 1^−4^ and alignment coverage < 80% of query gene length. Pearson correlation of iModulon gene coefficients between PCC 7942 and MED4 was used to identify conserved transcriptional modules (r > 0.20) (42); a deliberately inclusive threshold accounting for expected attenuation of cross-species correlation due to extensive gene content divergence.

### 5.7. Motif finding in transcriptional modules

DNA-binding sites for cyanobacterial TFs NtcA, SphR (PhoB), and LexA were obtained from RegPrecise (76). RpaA and RpaB motifs were built from ChIP-seq binding sites (6). Motifs were compiled using MEME v5.4.1 (77). Transcription units were defined using BioCyc operon annotation (78). Promoter regions were defined as 300 bp upstream of the first gene in each operon using pyranges v1.1.7 (79). FIMO v5.4.1 (80) was used to search for motifs in promoter regions, where the best-scoring motif for a given genomic region was retained (**Supplemental Data 6**).

### 5.8. Statistical testing

**Differential iModulon activities (DiMAs)** were computed as described previously (20). Briefly, replicate-average iModulon activities were compared between extended light periods and darkness, and between extended dark periods and light. The absolute differences were evaluated against a fitted log-normal distribution of all pairwise activity differences, with p-values corrected for multiple testing using the Benjamini–Hochberg method. iModulons were considered differentially activated if the p-adjusted value was less than 0.05 for both time points in each extended period of light or dark.

**Differential gene expression analysis** was performed with DESeq2 v1.46 (43). Gene expression was considered differentially expressed if the *p-adjusted* value returned by DESeq2 was less than 0.05 for both extended light and extended dark comparisons.

**Cosinor regression** was performed to assess circadian rhythmicity in iModulon activity and gene expression. A single-component cosinor model was fit to each time series, and statistical significance was evaluated by F-test against a null model, as implemented in CosinorPy (81). *P-values* were then adjusted for multiple testing using the Benjamini–Hochberg correction to determine significantly rhythmic genes (*p-adjusted* < 0.01) and rhythmic iModulon (*p-adjusted* < 1e^−3^).

## Supporting information

Supplemental Data 1 rnaseq_datasets

Supplemental Data 8 circadian_light_perturbed

Supplemental Data 7 circadian_gene_imodulon_fit

Supplemental Data 6 MED4_regulator_dna_motifs

Supplemental Data 5 MED4_PCC7942_BBH_iModulon_homology

Supplemental Data 4 pcc7942_imodulon_data

Supplemental Data 3 med4_imodulon_enrichment

Supplemental Data 2 med4_imodulon_data

## Acknowledgements

The research described in this paper was supported by the NW-BRaVE for Biopreparedness project funded by the U. S. Department of Energy (DOE), Office of Science, Office of Biological and Environmental Research, under FWP 81832. Z.J. is grateful for support from the Pacific Northwest National Laboratory (PNNL)-Washington State University (WSU) Distinguished Graduate Research Program (DGRP) Fellowship. Figures in this manuscript were created using BioRender.

## Supplemental Material

**Supplemental Results (**MED4_imodulon_manuscript_Supplemental_ZJ_022026.docx)

**SD1 (sd_1_rnaseq_dataset.xlsx):** RNA-seq dataset

**SD2 (sd_2_med4_imodulon_data.xlsx):** MED4 iModulons

**SD3 (sd_3_med4_imodulon_enrichments.xlsx):** Enrichments in MED4 iModulons

**SD4 (sd_4_pcc7942_imodulon)_data.xlsx):** PCC 7942 iModulons

**SD5 (sd_5_MED4_PCC7942_BBH_imodulon_homology.xlsx):** BBH iModulon homologs

**SD6 (sd_6_MED4_regulator_dna_motifs.xlsx):** Regulator DNA-binding motif hits

**SD7 (sd_7_circadian_gene_imodulon_fit.xlsx):** Circadian fit MED4 iModulons & Genes

**SD8 (sd_8_circadian_light_perturbed.xlsx):** Differential iModulon activity & gene expression in light perturbations

## Data availability

### RNA-seq Datasets

- Circadian Dataset: Primary RNA-seq data are openly accessible for download at the Gene Expression Omnibus (GEO) community repository under the data accession GSE314951
- Perturbation Dataset: [GEO accession: XXXXX]
- P-SSP7 infection Dataset [GEO accession: XXXXX]

### Code

- Zenodo: [BRaVE Zenodo: https://www.pnnl.gov/projects/nw-brave]
- GitHub: [Need to transfer from GitLab to https://github.com/NWBRaVE]
  - Current GitLab: [https://gitlab.pnnl.gov/nwbrave/thrust0/med4_ica_model]

### Author Contributions

**P.B., W.Q., M.S.C.:** Funding acquisition, Project management. **P.B., S.F., W.Q., M.S.C.:** Research & experimental design. **P.B., X.L., N.C.S., M.R.G.:** Performed experimental work to generate RNA-seq data. **Z.J.:** Computational analysis, interpretation of results, and data visualization. **Z.J., P.B.:** Writing — original draft. **Z.J., P.B., L.A.:** Data curation and data packing for public release. **Z.J., P.B., W.Q., T.Z., J.R.:** Writing — review & editing.

## Notes

https://www.pnnl.gov/projects/nw-brave

https://gitlab.pnnl.gov/nwbrave/thrust0/med4_ica_model

https://github.com/NWBRaVE]

https://www.ncbi.nlm.nih.gov/bioproject/PRJNA1393382

https://www.ncbi.nlm.nih.gov/geo/query/acc.cgi?acc=GSE314951

### Competing Interest Statement

The authors have declared no competing interest.

